# Independent recruitment of PRC1 and PRC2 by human XIST

**DOI:** 10.1101/2020.09.21.305904

**Authors:** Thomas Dixon-McDougall, Carolyn J. Brown

## Abstract

XIST establishes inactivation across its chromosome of origin, even when expressed from autosomal transgenes. To identify the regions of human XIST essential for recruiting heterochromatic marks we generated a series of overlapping deletions in an autosomal inducible XIST transgene. We examined the ability of each construct to enrich its unified XIST territory with the histone marks established by PRC1 and PRC2 as well as the heterochromatin factors MacroH2A and SMCHD1. PRC1 recruitment required four distinct regions of XIST, and these were completely distinct from the two domains crucial for PRC2 recruitment. Both the domains required and the impact of inhibitors suggest that PRC1 is required for SMCHD1 while PRC2 function is necessary for MacroH2A recruitment, although incomplete overlap of regions implicates a role for additional factors. The independence of the PRC1/PRC2 pathways, yet important of all regions tested, demonstrate both modularity and cooperativity across the XIST lncRNA.

**Author Summary:** XIST functions as a long, non-protein coding, RNA to initiate various pathways for the silencing of one of the two X chromosomes in female placental mammals. CRISPR-directed mutations of an inducible human XIST construct in somatic cells allowed us to discover which regions of the RNA are required for chromatin modification and protein recruitment. This was the first large-scale dissection of human XIST domains, and every function assessed was dependent on multiple regions of XIST, suggesting considerable interactions between domains of XIST. We observed similarities, but also differences, with the domains previously identified in mouse Xist and demonstrated the presence of independent pathways for chromosome reorganization in humans as well as ascribing new functionality to regions of XIST. The ability of XIST to inactivate large sections of chromosomes from which it is expressed makes it both an exciting potential therapeutic for chromosome number abnormalities as well as a paradigm for how non-coding RNA genes are able to regulate cellular biology.

## Introduction

XIST was one of the first long noncoding RNAs (lncRNA) to be identified (1), and its role in establishing the complex heterochromatin of the inactive X chromosome (Xi) continues to yield new insights into the process of X-chromosome inactivation (XCI) and the functionality of lncRNAs. Most studies of Xist have been conducted in mice; yet we know there are substantial differences in both the RNA and the XCI process between species (2). Human and mouse XIST/Xist show a similar exon/intron structure, including two major splicing isoforms, and ∼67% sequence conservation across the 15-19 kb lncRNAs (3). Both contain a series of tandem repeats (labelled A-F), of which only the A repeat is highly conserved in sequence and size across eutheria (Minks, Baldry, Yang, Cotton, & Brown, 2013) (4). The F and E repeats, which have limited tandem repeat structure, are also retained in both species. The most divergent repeat regions are B and C – the regions reported in mouse to be critical for recruitment of the polycomb repressive complexes PRC1 and PRC2 (5). The human B repeat region is split into Bh and B repeats, and human has only a fraction of one C repeat, while mouse has 14 copies. Additionally, in humans there are more copies of the D repeats than in mouse (3,6,7). Thus, the study of human XIST adds an important comparator to our growing understanding of mouse Xist functionality. In addition, roles for XIST variation in human disease will require an understanding of the functional domains of XIST, as will reducing the size of XIST to enable its potential as a ‘chromosome therapeutic’ to silence trisomic chromosomes (8, 9).

As inactivation normally occurs early in mammalian development, many alternative model systems for the study of XIST and XCI have been described, most of them utilizing mouse models (10). Model systems to study human XIST are more limited. The second X chromosome in human female ESCs is generally a mix of inactivated, eroded and active states (11). Rare *in vivo* studies in humans emphasize differences between human and mouse XCI, with both human X chromosomes expressing XIST prior to random silencing (12). In contrast, mouse has initial paternal inactivation that is reactivated prior to subsequent random inactivation in embryonic tissues, as well as regulatory elements including the antisense Tsix are not conserved outside of rodents (13–15). Transgenic human XIST has been used in induced pluripotent stem cells to generate very complete autosomal silencing (8), and also shown to induce many features of XCI in human somatic cells (16, 17). Recently, CRISPR-induced deletions in aneuploid 293 cells have shown the importance of some human regions for maintenance of XCI, and particularly for splicing (18).

Both the human and mouse Xi are seen to acquire repressive epigenetic alterations, with H3K27me3 and H2AK119ub1 (ubH2A) the result of PRC2 and PRC1 activity respectively (reviewed in 19). SMCHD1 and MacroH2A are acquired later in mouse differentiation, with SMCHD1 recruitment in mouse dependent on PRC1 (20–22). There has been considerable controversy over the order of PRC1 and PRC2 recruitment and the nature of the elements and intermediary proteins involved (reviewed in 23–25). A series of proteomic analyses identified numerous Xist-interacting proteins, including HNRNPK, which is now believed to bind the B/C repeats and recruit PRC1 that then enables PRC2 recruitment (5,26–28). Over 200 Xist-binding proteins include other hnRNPs, splicing factors, RNA modifying factors and chromatin modifying factors have been identified through various screening approaches (26,29,30).

We previously examined XIST induced from nine different integration sites, and while all sites recruited some UbH2A and suppressed Cot1 RNA expression, the recruitment of H3K27me3, macroH2A and SMCHD1 was variable between integration sites and did not appear to be dependent on each other (17). To follow up investigating the ability of XIST to modify its surrounding chromatin we chose the 8p integration site for its reliable recruitment of heterochromatic marks while also being distinct from the unique combinations of DNA elements of the X chromosome.

In this study we dissected the modularity of human XIST by comparing the ability of clones with deletions tiled across the XIST RNA in our inducible construct to recruit H3K27me3, UbH2A, macroH2a and SMCHD1. Recruitment of each mark required multiple distinct regions, and the regions involved in recruiting H3K27me3 were distinct from ubH2A. The observed independence of PRC1 and PRC2 recruitment in our inducible system was reinforced by treatment with a PRC1 inhibitor impacting only PRC1 and SMCHD1, while inhibition of PRC2 did not impact ubH2A, only H3K27me3, as well as MacroH2A. While the need for PRC1 for SMCHD1 recruitment is consistent with results from mouse development, the independence of PRC1 and PRC2 recruitment demonstrates recruitment by human XIST in the HT1080 cells is clearly reliant upon different regions of the lncRNA.

## Results

### Induction of autosomal XIST establishes heterochromatin within 5 days

Induction of the 8p human XIST cDNA transgene in the HT1080 fibrosarcoma cell line has been shown to recruit the PRC1/2 established histone marks, ubH2A and H3K27me3 (17). The XIST cDNA sequence remained transcriptionally inactive in uninduced cells due to binding of TetR to elements within the CMV promoter sequence (Figure 1A). Treatment of the cells with 1µg/ml doxycycline (dox) for five days (5ddox) resulted in the XIST RNA forming a discrete unified domain that was frequently visibly enriched for heterochromatin marks such as H3K27me3 using immunofluorescence and fluorescent *in situ* hybridization (IF-FISH) (Figure 1B). To capture cell heterogeneity and overcome operator bias we developed a method of quantifying the extent of enrichment in individual cells. We calculated the relative fluorescent intensity and variation of a given heterochromatin mark at the site of XIST RNA in the nucleus compared to a cross section of the nucleus. In each cell the relative enrichment of the heterochromatin mark at the XIST RNA locus was expressed as a z-score, calculated relative to the variability of heterochromatin density across the nucleus and XIST RNA signal (Figure 1C). A z-score of 1 indicated that the median fluorescent intensity of a heterochromatin mark at the XIST RNA cloud was a full standard deviation above the average across the nucleus. In mouse embryonic stem cells (ESCs), enrichment of ubH2A and H3K27me3 occurs rapidly upon Xist expression (31); however, in these human differentiated cells we considered that recruitment might be slower given that epigenetic states are less plastic than in embryonic cells. Thus, to determine if marks were fully established by 5 days of induction, we calculated the relative enrichment of the PRC-associated marks H3K27me3 and ubH2A at the XIST RNA cloud following 3,5 or 10 days of dox (Figure 1D). Both marks were still accumulating following 3 days of XIST induction, with the median z-score for ubH2A enrichment 0.96 and H3K27me3 slightly higher at 1.48 across each population of 60 cells. By day 5 of dox induction the median z-score of the enrichment of H3K27me3 increased to 2.6, and remained constant after 10 days dox (z-score = 2.7). The enrichment of ubH2A increased more dramatically by day 5, with a median z-score of 4.1; however, enrichment then declined by day 10 to z-score = 1.8, which was still more than double the level at day 3 (Figure 1D). Changes were not attributable to differing levels of XIST RNA, as qPCR demonstrated that XIST RNA levels were not statistically different between 2 and 5 days of dox induction (Rq 2ddox = 0.68 relative to 5ddox, p = 0.24, supplementary figure 1).

**Figure 1:**
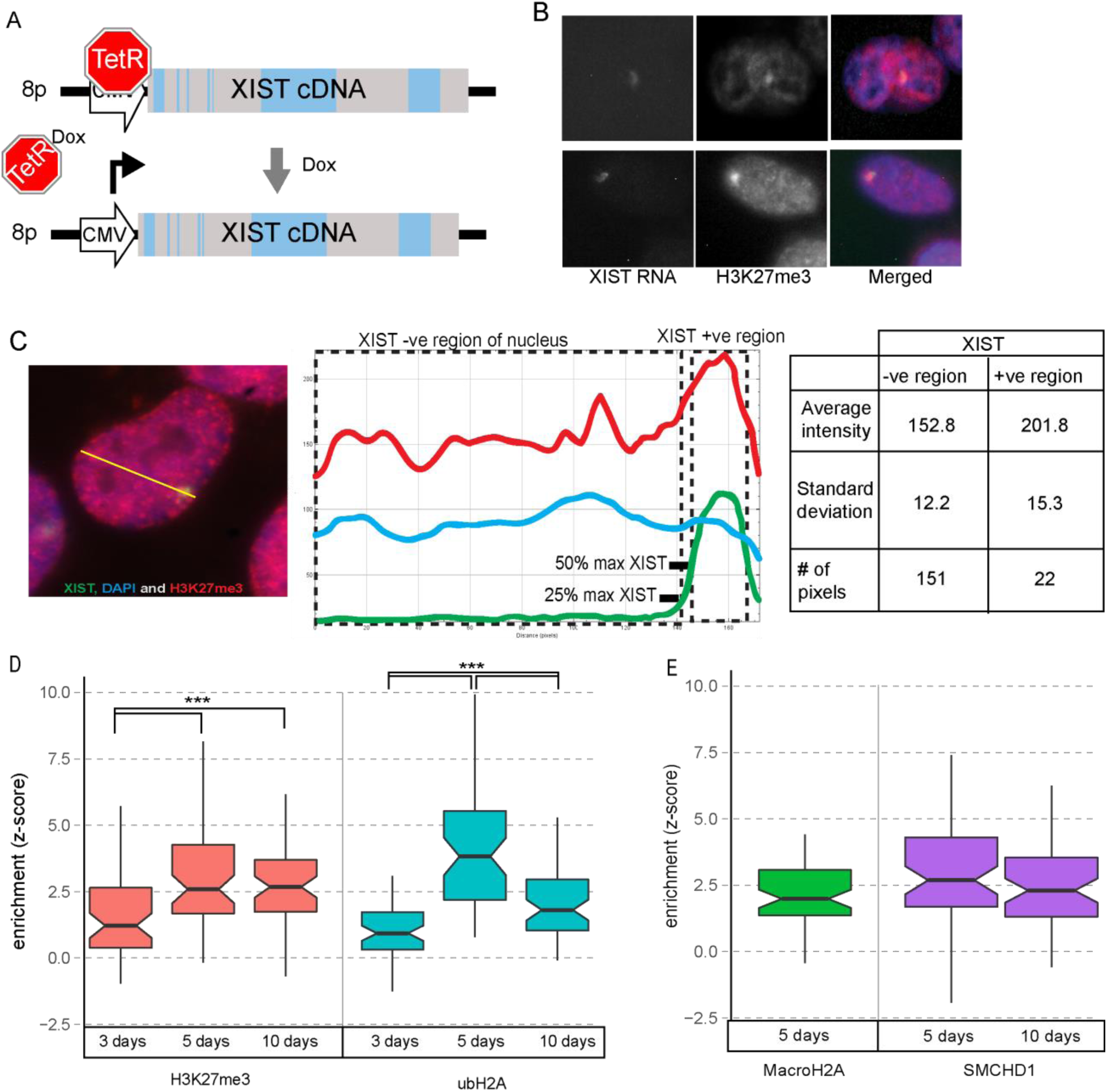
Autosomally expressed XIST RNA maximally enriches its surrounding chromatin with heterochromatin by day 5. A) Diagram of XIST cDNA construct under control of a Doxycycline (dox) inducible CMV promoter integrated into chromosome 8p of HT1080 cells. B) Example images of IF-FISH labelled HT1080 cells expressing XIST RNA with visible H3K27me3 enrichment. C) Schematic of how the levels of a chromatin mark (e.g. H3K27me3) were measured at the XIST RNA cloud relative to the nuclear background. D) Enrichment (z-score) of H3K27me3 and ubH2A after different periods of XIST induction from chromosome 8p. E) Test of MacroH2A and SMCHD1 enrichment after five days of XIST induction as SMCHD1 after 10 days of induction. D-E) 60 cells were measured for each condition, with the centre denoting the median, notch indicating confidence interval and box extending from the 25th to 75th percentile of each population. Significance calculated using the Mann Whitney U test (*** p < 0.001).

In addition to examining the PRC1/2 recruited marks we wanted to examine the enrichment of SMCHD1 and macroH2A by XIST induction. Using the same procedure to quantify the relative distribution of these marks at the XIST RNA cloud we observed a clear enrichment of both marks (Figure 1E). The median z-score of MacroH2A enrichment across the population of HT1080 cells was 2.0, lower than the other marks examined at this time point. The median z-score of SMCHD1 was 2.7 (Figure 1F); however, as its enrichment had been associated with ubH2A in mice we were curious whether it would also decrease by day 10. The level of SMCHD1 enrichment was similar at day 10 (median z-score = 2.3) with the small decrease relative to its level at day 5 not reaching statistical significance (p = 0.054, Figure 1F).

### Repeat regions of XIST are not required for expression or localization of the lncRNA

The human and mouse XIST/Xist lncRNAs are large with functionality assigned to multiple of the mouse repeat regions. To assess the presence of functional domains within human XIST we generated a series of deletions, including not only the repeats but also non-repeat regions. Cell lines with partial XIST cDNA constructs were recombined into the 8p integration site (16): Exon 1 consisted of all but the 3’ ∼3.6kb of XIST and ΔPflMI had an internal ∼3.8kb including Repeat B and C removed (Figure 2A). In addition, an XIST deletion construct, ΔΔ, was created that removed both regions from the ΔPflMI and Exon 1 constructs. While these constructs provided a framework for examining large sections of XIST, a more complete dissection of XIST was desired to identify all potential regions crucial for the recruitment of both PRCs by XIST. Therefore, a series of nine additional deletion constructs were created by excising sections of the full length XIST construct integrated into chromosome 8p of the HT1080 cells using CRISPR, an approach that would also minimize potential genetic or epigenetic drift between clones (Figure 2B). These were designed with an emphasis placed on specifically excising the tandem repeat sequences separately from one another, although the human B and C repeats were too small and close to each other to be separately removed. The long non-repeat sequences of XIST were also removed individually, such that the entire XIST transgene would be interrogated for potential roles in XIST-mediated heterochromatin recruitment. The gRNAs selected to generate the various deletion constructs were designed to create small partially overlapping deletion sequences between adjacent deletions to maximize their resolution investigating XIST (Supplementary Table 1). Independently generated clonal cell lines were isolated for each deletion construct, and the region surrounding the excision of each confirmed through sequencing (Supplementary Table 2).

**Figure 2:**
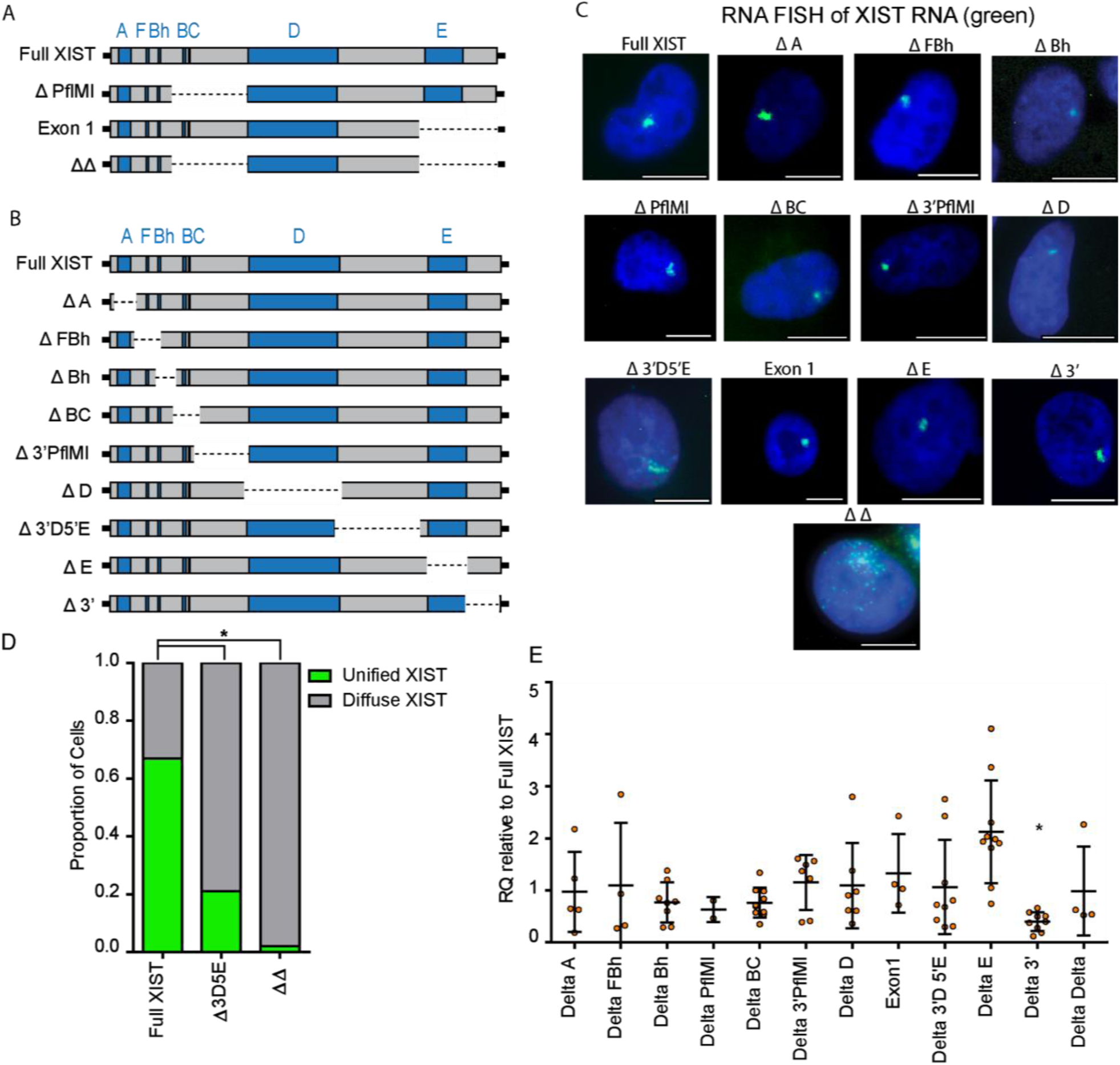
XIST deletion constructs used to investigate the regions of XIST crucial for chromatin remodelling. A) Illustration of the original full length and partial XIST constructs independently recombined into the FRT site on chromosome 8p of the HT1080 cell line. B) The extent of the XIST deletion constructs generated through the use of CRISPR technology from the Full XIST HT1080 cell line. C) FISH images illustrating the typical XIST RNA cloud (green) produced by each of the deletion constructs following 5ddox induction. D) Quantifying the proportion of XIST RNA clouds that were either clearly unified or punctate in appearance between the Δ3D5E and ΔΔ constructs in relation to Full XIST. Proportions were compared using Fisher exact test (* p < 0.01) E) Relative XIST RNA levels across numerous biological replicates for each of the deletion constructs. XIST RNA levels were calculated using the endogenous control gene PGK1 and were expressed relative to a Full XIST control and a t-test was used to calculate statistical significance, with the threshold adjusted for multiple testing (* p = 2.05×10^-5^).

Inducing XIST expression with doxycycline resulted in a unified XIST RNA signal that could be readily identified when examined by IF-FISH. Most of the XIST RNA signals for each of the deletion constructs as well as Full XIST were clearly unified into a single clearly distinct domain within the nuclei (Figure 2C). However, in the ΔΔ construct the absence of both the PflMI region and latter exons of XIST resulted in a clearly diffuse punctate XIST signal that was immediately distinguishable from Full XIST. The only other deletion construct that noticeably affected the distribution of XIST RNA in the nucleus was the Δ 3D5E construct lacking the non-repeat sequence of XIST between Repeats D and E. Quantification demonstrated that 67% of Full XIST RNA signals across 330 cells localized into one clearly unified XIST RNA signal with no observable gaps between regions of peak signal intensity. In contrast only 21% of XIST RNA signals in 380 Δ3D5E cells were clearly unified, and only 2% of the 271 ΔΔ cells examined had a unified XIST RNA signal (Figure 2D). While beyond the scope of this current research these results suggested that redundant and potentially additive elements involved in XIST RNA unification were present across distinct regions of XIST.

Finally, it was tested whether the various deletion constructs impacted the relative levels of XIST RNA induced by 5ddox. XIST RNA levels in the deletion constructs were compared relative to the levels found in Full XIST constructs (Figure 2E, Supplementary Table 3). Only the Δ 3’ construct was observed to be statistically different (p = 2.0×10^-5^), with less expression than Full XIST, suggesting that there may be some elements in that 3’ region that contribute to transcript stability. The lower transcript levels of the Δ 3’ region were not observed in the encompassing Exon 1 deletion construct (RQ = 1.32, p = 0.46, Figure 2E), suggesting that the effect was based on more complex mechanisms than simply the absence of stabilizing factors.

### Distinct regions of XIST crucial for PRC1 and PRC2 recruitment

Our foremost objective was to identify the regions of XIST that were crucial for recruitment of the two types of PRC-established marks, H3K27me3 and ubH2A. We first examined H3K27me3 across the HT1080 Δ constructs to identify which region(s) of *XIST* were essential for recruitment and/or activation of PRC2. H3K27me3 fluorescence at the Full *XIST* RNA cloud was noticeably enriched (median z-score = 2.59, sd = 1.70) which can be conceptualized as the average H3K27me3 intensity at XIST being roughly equal to the 99^th^ percentile of H3K27me3 fluorescent intensity measured in the nucleus (Figure 3A). Most of the deletion constructs showed comparable levels of H3K27me3 enrichment, indicating that over 80% of XIST could be removed without noticeably disrupting this process. Two regions of XIST were observed to be crucial for XIST to enrich the surrounding chromatin with H3K27me3, demonstrating a statistically significant difference from Full XIST levels of enrichment. The ΔFBh construct completely lacked H3K27me3 enrichment across its population of cells, with an average z-score of −0.147 (p = 1.11×10^-13^, Figure 3A and supplementary table 4). The overlapping deletion ΔBh showed an attenuated enrichment of H3K27me3 (median z-score = 0.787) that still differed significantly from Full XIST (p = 8.47×10^-07^, Figure 3A). The ΔA construct also had a statistically attenuated enrichment of H3K27me3 (1.59, p = 1.16×10^-4^); however the magnitude of this effect was clearly less extreme than for the adjacent ΔFBh construct, as ΔA and ΔFBh were strongly dissimilar (p = 1.49×10^-9^). Repeat E of XIST was identified as a second, entirely distinct, region of XIST indispensable for H3K27me3 enrichment, as the absence of this region in the ΔE construct resulted in no observable enrichment and a strong statistical difference from Full XIST (median z-score = −0.010, p = 2.97×10^-17^, Figure 3A). The role of Repeat E was supported by the similar scores seen for the overlapping deletions of Exon 1 and ΔΔ.

**Figure 3:**
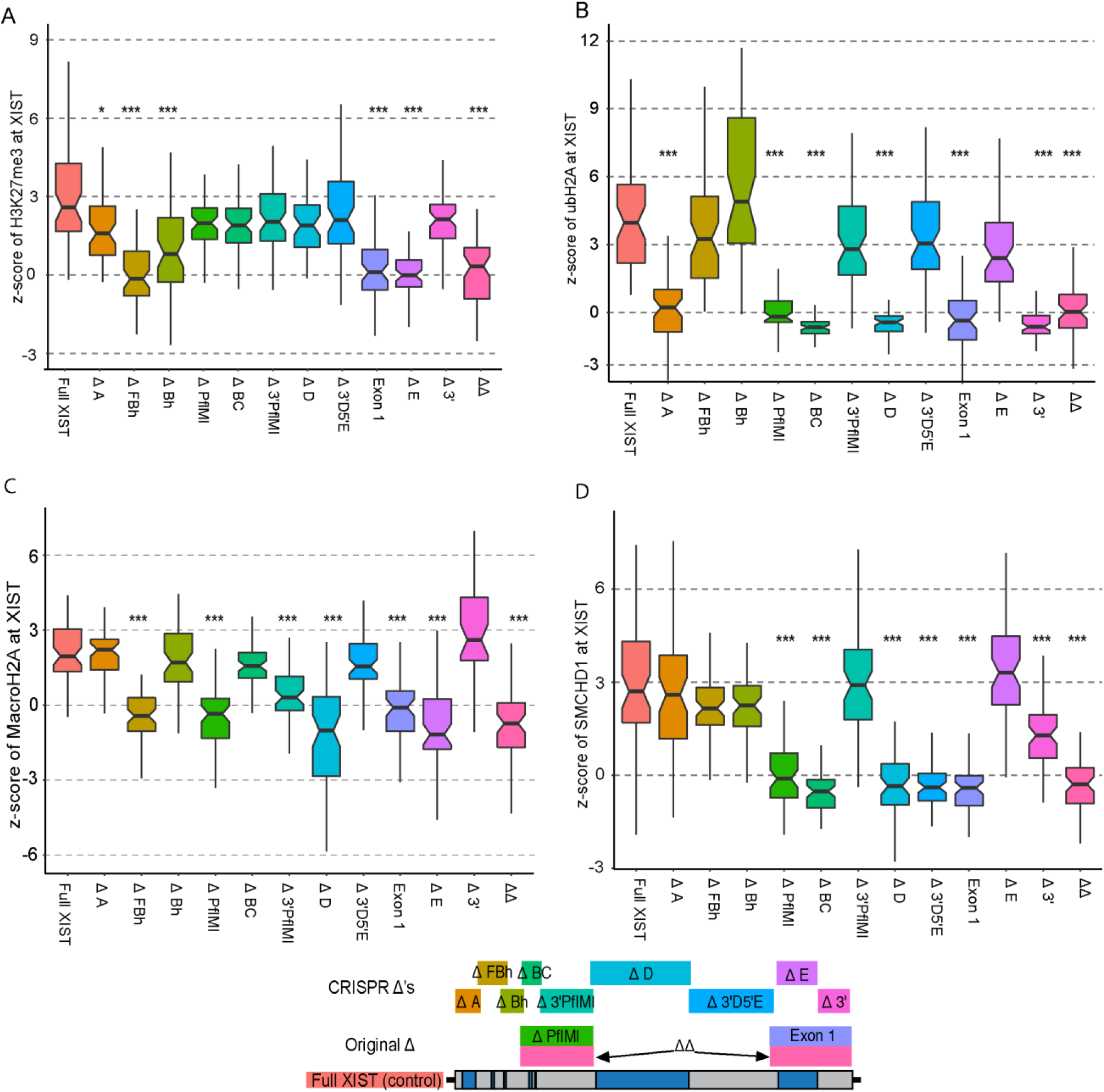
Multiple distinct regions of XIST are necessary for each type of chromatin modification. Each deletion construct induced for 5 days was tested for its relative enrichment (z-score) of one of the Xi associated heterochromatin marks (59-61 cells per construct). The XIST deletion constructs were tested for their ability to enrich their chromatin with A) H3K27me3, B) ubH2A, C) MacroH2A, D) SMCHD1. We were blinded to the identity of every cell analyzed in this study and only unblinded once all data had been collected into a single table. The population distribution of enrichment across the cells of each deletion construct were statistically compared to Full XIST using a Mann Whitney test with multiple testing correction (48 tests in total, * p-value < 1.04×10^-3^, ** p-value < 2.08×10^-4^, *** p-value < 2.08×10^-5^).

Previous work has indicated a linkage between PRC1 and PRC2 recruitment in mouse models for XCI, so we set out to investigate whether a similar link might exist for human XIST with the same regions involved in recruitment of both complexes. The deletion constructs were tested for their ability to enrich their surrounding nuclear territory with ubH2A to identify the crucial regions of XIST for PRC1 recruitment (Figure 3B and supplementary table 5). Full XIST showed greater relative enrichment of ubH2A than any other heterochromatin feature, although with a much greater standard deviation across the 60 cells (median z-score = 4.18 +/-3.18). The constructs that had disrupted H3K27me3 enrichment still showed strong ubH2A enrichment, as did constructs lacking the non-repeat regions on either side of Repeat D. Enrichment of ubH2A was found to be highly dependent upon the repeat regions in the first exon of XIST with ΔA, ΔBC and ΔD all showing no evidence of enrichment in the population of cells examined, and all strongly statistically different from Full XIST (median z-scores ≤ 0.223, p ≤2.84×10^-17^). In addition, the 3’ most deletion construct, Δ3’, also failed to enrich chromatin with ubH2A (median z-score = −0.635, p = 4.80×10^-21^) suggesting that this small ∼630nt region was also crucial for PRC1 activity and/or recruitment. It was unlikely that this effect was due to the lower transcript levels of the Δ3’ construct cell lines as the encompassing deletion of the Exon 1 constructs also failed to enrich for ubH2A (median z-score = −0.352) despite having XIST transcript levels slightly greater than Full XIST (Figure 2E). These results revealed four distinct regions of XIST crucial for the enrichment of ubH2A at the XIST RNA territory, none of which were crucial for PRC2 enrichment and *vice versa*.

### Overlap in regions of XIST crucial for PRC1/2 with those for SMCHD1 and MacroH2A

Given our observation that PRC1 and PRC2 were recruited by independent domains of XIST in our human cells, we wished to determine if either of the marks that are reported to be recruited later during mouse XCI would be dependent on similar domains. MacroH2A is a well-established Xi-associated mark, known to be recruited by Xist, but had not previously been associated with other components of the XCI pathway. By examining which regions of XIST are crucial for MacroH2A enrichment we sought to reveal additional hierarchies for chromatin remodelling during XCI. MacroH2A had a somewhat weaker relative enrichment compared to the other Xi-associated factors described here, with the Full XIST population having a median z-score of 1.96 (Figure 3C and supplementary table 6). As with the other XCI factors examined, numerous distinct regions of XIST were identified as crucial for MacroH2A enrichment. The deleted regions in the ΔFBh and ΔE constructs identified as crucial for H3K27me3 enrichment were also found to be crucial for MacroH2A enrichment (median z-score ≤ −0.410, p ≤ 6.08×10^-17^, Figure 3C). A third broad region encompassing Repeat D and the non-repeat region upstream of it (Δ3’PflMI) was found to be essential for MacroH2A enrichment (median z-score ≤ 0.313, p ≤ 3.06×10^-09^, Figure 3C). The results of this analysis led to the intriguing idea that MacroH2A may rely on PRC2 recruitment or activation by XIST, with additional factors located upstream and within Repeat D of XIST.

SMCHD1 recruitment by Xist has been reported to require mouse repeats B and C to recruit PRC1 (Jansz). Since we found ubH2A enrichment in humans to also associate with Repeats B and C, despite the divergence in size and position of these repeats between species, we questioned whether the association between PRC1 and SMCHD1 might also exist in humans. We compared the enrichment of SMCHD1 at the XIST RNA cloud in the deletion constructs relative to the Full XIST control as described in the previous sections (Figure 3D and supplementary table 7). The enrichment of SMCHD1 at Full XIST following 5ddox induction was found to be 2.704, and the three 5’ most Δ constructs had similar levels of enrichment (median z-score ≥ 2.16), indicating that those regions that had been associated with H3K27me3 enrichment were dispensable for SMCHD1 enrichment. The ΔBC and ΔD constructs that had been unable to enrich ubH2A were also found to be incapable of enriching SMCHD1 (median z-score ≤ −0.350). The Delta 3’ terminal region that had been incapable of recruiting ubH2A, however, showed a weak but still observable enrichment of SMCHD1 (median z-score = 1.291), suggesting that the region contributed to SMCHD1 enrichment without being essential. Finally, the non-repeat region of XIST between Repeat D and E was also found to be crucial for SMCHD1 enrichment as the Δ3D5E deletion failed to enrich its nuclear territory with the factor and differed significantly compared to Full XIST (median z-score =-0.386, p = 6.35×10^-17^). Thus, while the B-C-D and 3’ regions were required for both ubH2A and SMCHD1, additional elements were involved in the recruitment of each by XIST.

### Inhibititors confirm independence of PRC1/SMCHD1 from PRC2/MacroH2A

Perhaps the most surprising insight from our analysis of the XIST Δ constructs was the apparent independence of recruitment of PRC1 and PRC2 by XIST in human somatic cells. To test this independence we treated the HT1080 cells with either the PRC2 small molecule inhibitor GSK343 or the PRC1 small molecule inhibitor PRT4165. Both of these inhibitors only affect the enzymatic activity of PRC2 or PRC1 rather than damaging the complexes and both inhibitors were demonstrated in numerous previous studies to be highly specific for their target enzyme (32, 33). Using these inhibitors would also test our hypotheses that SMCHD1 and MacroH2A were reliant on PRC1 and PRC2, respectively. For the 5 days that the cells were undergoing dox treatment to induce XIST expression we concurrently treated the cells with the inhibitors at levels reported to be effective at inhibiting either PRC2 or PRC1, but that did not cause visible signs of stress to the cells or disrupting the XIST RNA cloud.

Chemical inhibition of PRC2 with 5µM GSK343 resulted in a clear disruption of H3K27me3 enrichment compared to the uninhibited control (median z-score = 0.162, p =7.24E-16 Figure 4A, supplementary table 8). These results suggested that the treatment was effective at disrupting the enrichment of H3K27me3 by XIST. We then examined whether inhibition of PRC2 affected PRC1 mediated ubH2A enrichment by XIST. In keeping with the expectation of independence based on the examination of the Δ constructs, we observed that ubH2A was still strongly enriched at the XIST RNA cloud even when PRC2 was inhibited (median z-score = 3.053, p = 0.22, Figure 4A). We also examined whether disruption of PRC2 affected MacroH2A, as the overlapping regions of XIST crucial for each suggested a potential functional link. We observed that inhibition of H3K27me3 completely disrupted MacroH2A enrichment (median z-score = 0.224) and this was strongly statistically different from the uninhibited induced Full XIST control (p = 1.30×10^-10^, Figure 4A). These observations led to the conclusion that the activity of PRC2 at the XIST RNA cloud was an essential step for the enrichment of MacroH2A.

**Figure 4:**
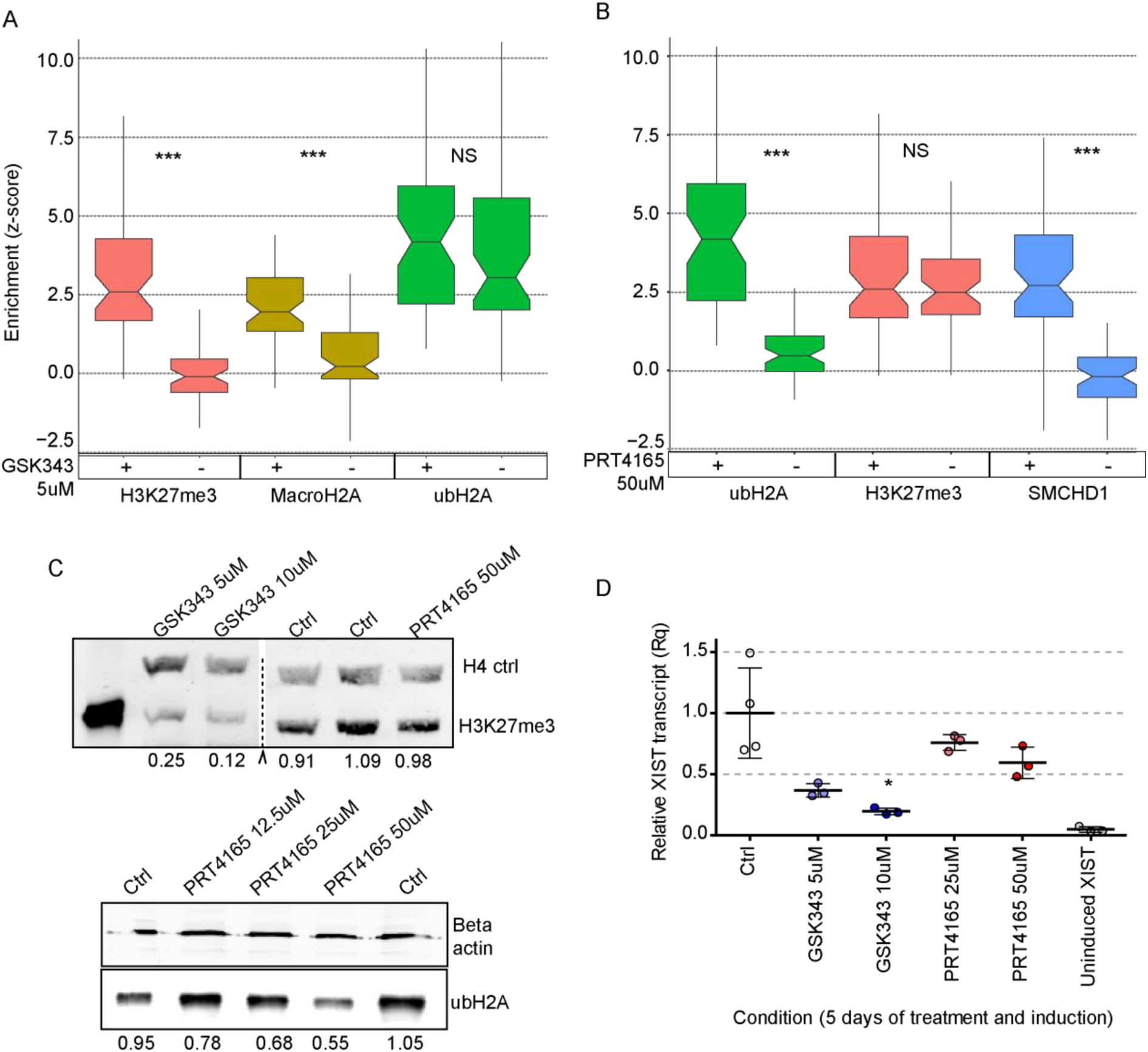
The two polycomb complexes operate independently to promote further heterochromatin formation. The effect of A) PRC2 inhibition with GSK343 or B) PRC1 inhibition with PRT4165 on the relative enrichment of H3K27me3 (pink), MacroH2A (yellow), SMCHD1 (blue) and ubH2A (green) at the XIST RNA cloud relative to the average level in the nucleus across a population of 60-61 cells. We were blinded to all cells and chromatin marks identity until after all data and calculations were completed. Statistical significance of the effect of each inhibitor on a chromatin mark was calculated using the Mann Whitney test with adjusted p value (*** p < 1×10^-6^). C) Western blotting images demonstrating the levels of ubH2A and H3K27me3 after cells had undergone chemical inhibition with either PRT4165 or GSK343. The dotted line denotes where several bands from an unrelated set of tests were spliced out digitally. The numbers underneath each row of bands denotes the relative concentration of the chromatin mark being tested compared to the control populations. D) Relative XIST RNA levels for each the inhibitor concentrations relative to control 5ddox. The dots denote independent biological replicates and statistical significance was calculated using a t-test (* p < 0.05).

Inhibition of PRC1 with 50µM of PRT4165 had the anticipated effect of preventing the XIST RNA cloud from becoming enriched with ubH2A relative to the nuclear background (median z-score = 0.469, significance relative to Full XIST p = 5.28×10^-18^, Figure 4B, supplementary table 8). Inhibition of PRC1 and the lack of ubH2A at the XIST RNA cloud did not affect the ability of XIST to enrich H3K27me3 (median z-score = 2.483), further supporting the independence of PRC1 and PRC2 recruitment by XIST (Figure 4B). PRC1 activity, however, was found to be crucial for SMCHD1 enrichment, as PRC1 inhibition resulted in a significant loss of SMCHD1 enrichment at the XIST RNA cloud (median z-score = −0.196, p = 5.28×10^-18^, Figure 4B). Therefore, a role for ubH2A enrichment for the recruitment of SMCHD1 to the XIST RNA territory may have been conserved between mice and humans.

Neither inhibitor completely removed its affected mark throughout the nucleus, allowing our continued use of the quantitation approach outlined in Figure 1C. We quantified the impact of the inhibitors by performing western blotting for H3K27me3 and ubH2A in the 8p Full XIST cells after 5 days of inhibition treatment and dox were compared to cells after 5 days of dox induction without inhibitors. Following 5 days of inhibition with GSK343 there was 25% the relative amount of H3K27me3 remaining in the cells compared to the control populations (Figure 4C). Following 5 days of PRC1 inhibition with PRT4165 roughly only 55% of ubH2A remained in the HT1080 cells relative to cells without the inhibitor. The PRC1 inhibition did not produce a statistically significant effect on XIST RNA levels compared to the uninhibited XIST transcript levels (Figure 4D). The PRC2 inhibitor by contrast produced a clear dosed-dependant decrease in XIST transcript levels (Figure 4D). The relative levels of XIST following PRC2 inhibition with the 5µM concentrations used in these experiments were comparable to those seen in the Δ3’ cell lines, and coupled with the clear enrichment of ubH2A in both cases it seemed that this lower expression level did not cause an obvious disruption to XIST function.

## Discussion

LncRNAs have been suggested to function as modular scaffolds for protein binding and the internal repeats of XIST/Xist have exemplified the concept (34, 35). In this study we sought to characterize domains of the human XIST, focussing not only on the repeat regions, but also interrogating the non-repetitive regions of the lncRNA. We generated 13 deletions that removed from 800 bp to over 3 kb of the cDNA. For each deletion we characterized the expression and localization of XIST as well as the ability to recruit H3K27me3, H2Aub, SMCHD1 and MacroH2A. As summarized in Figure 5, every feature required multiple regions of XIST to become enriched upon the XIST-coated chromatin, and there was no single critical region that was necessary for all features. While repeat-containing regions were crucial for the features analyzed it is worth noting that the traditionally less studied non-repeat regions were also found to be critical in our system.

**Figure 5:**
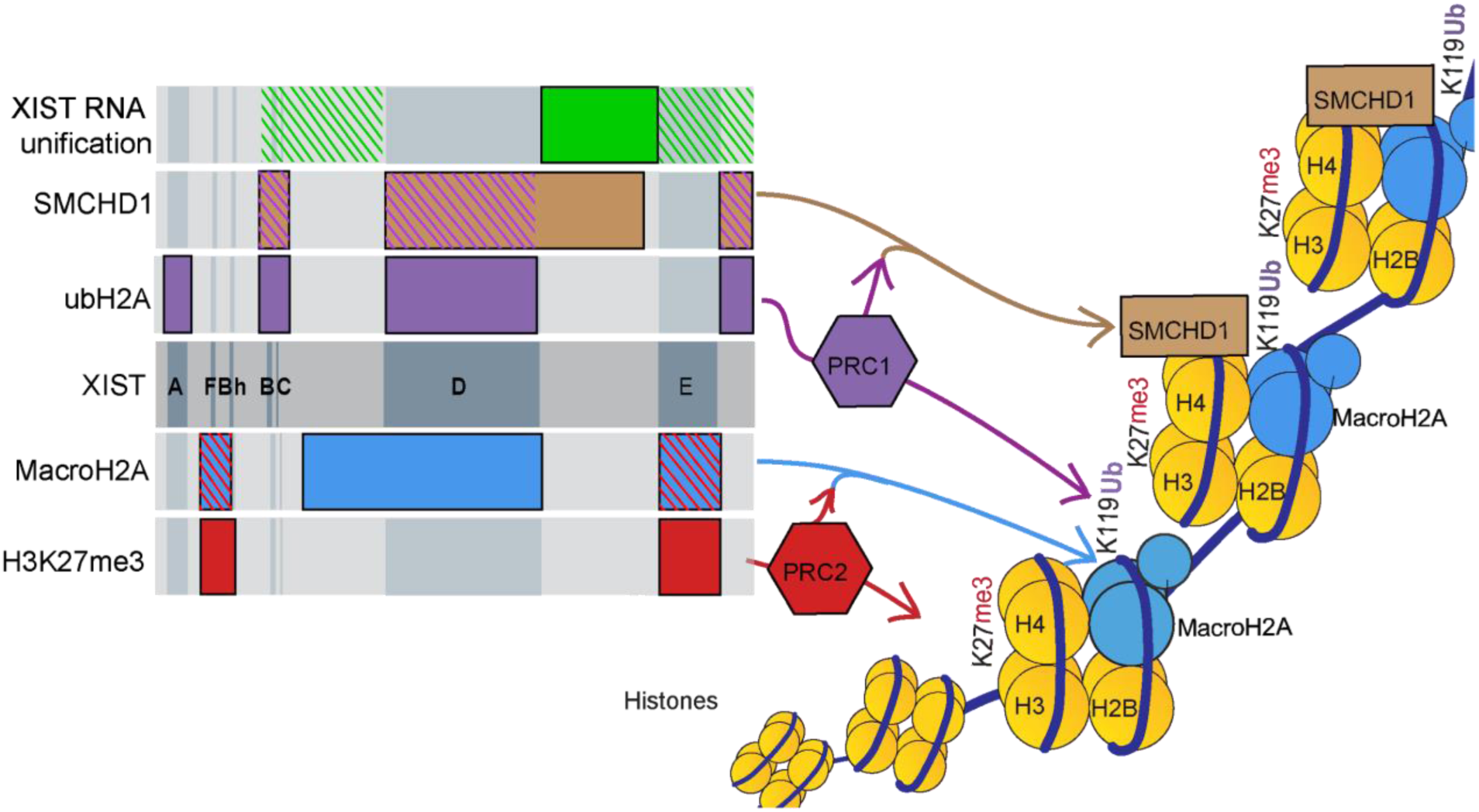
Summary of the regions and pathways proposed to be crucial for XIST and PRC mediated chromatin remodelling. PRC2 (red) and PRC1 (purple) were identified to operate through entirely independent regions of XIST and to not affect each other in this context. PRC2 catalytic activity was crucial for MacroH2A (blue) enrichment while PRC1 catalytic activity was critical for SMCHD1 (brown) enrichment. Regions of XIST identified as important for the recruitment of these chromatin features were marked in the relevant solid colour, while dashed lines denoted the PRC dependent regions that overlapped with the late chromatin marks. XIST RNA unification (green) was found to be affected by the loss of a large internal non-repeat sequence (solid colour) as well as two redundant regions of XIST (dashed lines).

Each feature required two or three regions of XIST separated by kilobases of RNA, that when independently deleted had no impact upon recruitment of the feature. Previous work had proposed that long-range interactions were endemic across XIST through the use of RNA duplex mapping approaches (34, 36). One could envision specific RNA secondary structures formed through long range interactions that may allow for complexes to bind to XIST, or distinct protein interactions with different RNA regions that are required to assemble essential protein complexes. While not exclusive of each other, the latter explanation may fit with evidence of five hubs of prolific protein binding, around the A repeats, F repeats, B-C-D repeats, E repeats and 3’ end of XIST (34). Studies in mouse models have argued that the multivalency of the E repeat region allows homo- and hetero-typic interactions with partial redundancy that function in promoting phase separation and formation of the inactive X compartment (37, 38). Such aggregation might be impacted by overall density of protein binding along the RNA.

Strikingly, our results showed that recruitment of the marks established by PRC1 and 2 relied upon entirely distinct domains in the human somatic cells with induced XIST expression. Similar to mouse, PRC1 recruitment required the BC region, which is considerably smaller in humans with only part of a single C repeat and a separation of the B into two clusters of repeats (3). The D repeat region was also required for human PRC1 recruitment, consistent with structural analysis suggesting a large protein-interaction domain containing from B-C to beyond the D repeats in exon 1, as well as HNRNPK binding to the human D repeats (34). Thus the B-C-D region may recruit PRC1, likely through HNRNPK, with weighting in humans towards the requirement including the D repeat region as B-C is smaller than in mouse. We identified two additional regions that were also essential for PRC1 recruitment - the A repeats, and the 3’ end of XIST. Again, in mouse models, it has been shown that silencing is necessary for the spread of UbH2A into genes (31), and it is possible that we only detect UbH2A when it has spread beyond initial deposition. The A repeats are also critical for binding of RBM15 for m6A deposition at the 5’ and extreme 3’ ends of XIST (34), so it is possible that modification of the RNA affects the ability of XIST to bind PRC1 (26,39,40).

The recruitment of H3K27me3 was impacted by only two regions - the F repeat region and the E repeat region. These regions do not overlap the B-C region seen to be necessary for murine PRC1 recruitment, nor the regions discussed above as being necessary for human PRC1 recruitment. Many regions and pathways have been reported to be responsible for recruitment of PRC2 in mouse (2), including repeats F and B for JARID2 recruitment for H3K27me3 (41) and a previous RNA immunoprecipitation study with human XIST found EZH2 and SUZ12 bound repeat E (42). Another consideration is whether the different regions of XIST modulate unique aspects of PRC2 recruitment versus activity, potentially as a result of the differing effects of the various core factors (SUZ12) and cofactors (e.g. JARID2) involved.

SMCHD1 recruitment overlapped regions required for PRC1, and was also inhibited by PRT4165, a specific PRC1 inhibitor, which is consistent with mouse studies showing SMCHD1 recruitment requires UbH2A (20). In our system, SMCHD1 also required the non-repetitive region at the end of exon 1 that has been seen to form numerous duplexes with the BC region of XIST. SMCHD1 did not require the A repeat region, perhaps reflecting that the silencing required to allow detectable spread of UbH2A is not required for the recruitment of SMCHD1 to the XIST-coated chromosome.

There was also overlap between the regions of XIST required for H3K27me3 and MacroH2A recruitment, and inhibition with GSK343 substantially reduced the recruitment of both to the XIST-coated domain, indicating that MacroH2A recruitment was dependent upon the catalytic activity of PRC2. The discovery of this novel association has yet to be investigated in mouse models, though both have been observed to be lost when Xist is deleted (43, 44). At time of writing the mechanisms underlying the connection between PRC2 and MacroH2A during XCI are unknown. MacroH2A is enriched at H3K27me3 enriched chromatin throughout the genome (reviewed in 45), but appears to be established independently (46). It remains to be determined whether MacroH2A directly interacts with XIST; however, the lack of its presence in the interactome screens may implicate intermediate factors, which perhaps bind upstream of the D repeat (26,29,39,47).

Overall, we observe similarities and differences from previous studies of the functionality of Xist/XIST. Many of these have utilized mouse ES cells, and thus our study differs in both developmental timing as well as the species being investigated. The difference between mouse and human sequences are discussed above; however, another important difference between humans and mice is the limited developmental window for Xist function, while we have seen that XIST can induce many features of XCI in human somatic cells, although the cells studied here are cancer-derived and thus may have a less restrictive chromatin state. The differences between the broad contexts of human and mouse XCI are extensive and have recently been well reviewed (48). A few notable differences include the relatively unique occurrence of imprinted Xist expression, differentiation-dependent protein binding and the more extensive inactivation of X-linked genes in mice relative to humans (15, 49).

In our system XIST is induced from a viral-derived inducible promoter, with the promoter inducing expression to approximately the same level as seen for endogenous XIST, an important consideration given the myriad and cooperative binding of proteins to XIST. Previous reports suggested that repeats D and F (also A??) are involved in expression of XIST (50–52), which would not impact our promoter. In fact only the specific removal of the 3’ non-repeat region of XIST was found to decrease transcript levels in our study. The unification of the XIST RNA transcripts themselves into a single domain was found to be generally very stable, with only the synergistic deletion of both >3kb regions of the ΔΔ construct and the deletion of the large non-repeat region spanning between repeat D and E noticeably affecting RNA unification. The non-repeat region spanning D to E was also found to be crucial for SMCHD1 enrichment through ubH2A independent mechanisms. SMCHD1 has been associated with the compartmentalization of the Xi, thus it is possible that the two processes of unification and SMCHD1 enrichment may be critically facilitated by this region of XIST as shown in summary Figure 5 (53, 54).

We only examined 4 chromatin marks in a somatic cell model, yet we found that every region of XIST we examined was critical for some aspect of XCI. Thus, despite the size of XIST, and the only limited conservation with mouse Xist, including deviation in extent of tandem repeats, the lncRNA seems highly adapted to function in multiple independent pathways. While our results contribute to the concept of modularity of lncRNAs, they also emphasize the need to shift the paradigm of the functional domain of a lncRNAs away from being single linear sequences towards being based on secondary and tertiary arrangements. Future studies and research examining how to regulate these complex and long range interactions between regions of XIST and other non-coding RNAs seem likely to yield new breakthroughs in the field of RNA biology and potential insights into the utility of XIST as a therapeutic for chromosome abnormality disorders such as trisomy 21.

## Materials and Methods

### Cell culturing protocols for the HT1080 cell lines

Dox-inducible XIST cDNA constructs in autosomes of the male fibrosarcoma cell line, HT1080, were generated as described by Kelsey et al and Chow et al. In brief, inducible XIST cDNA constructs (Delta PflMI, Exon 1 or Delta Delta; Figure 1) were integrated into an 8p FRT site in HT1080 cells previously transfected with pcDNA6/TR for Tet-Repressor protein expression. The full XIST cDNA (Full XIST) was previously described, and corresponded to the short-isoform of XIST, the mouse homolog of which has also been reported to be fully functional (55). The integrated XIST cDNA constructs were controlled by a CMV promoter, that was blocked by TetR and induced in the presence of 1ug/ml doxycycline. The HT1080 cells were grown at 37°C with 5% CO_2_ in DMEM supplemented with 10% Fetal Calf Serum (Sigma-Aldrich) by volume, 100 U/ml Penicillin-Streptomycin, non-essential amino acids and 2mM L-Glutamine. The chemical inhibitors GSK343 (Sigma-Aldrich) and PRT4165 (Sigma-Aldrich) were dissolved in DMSO and added to the DMEM media at the concentrations listed during the chemical inhibition assays. An equal volume of DMSO (without inhibitors) as was used in the inhibition treatments was added to the media of the HT1080 cells used as controls during the inhibition experiments to ensure that any differences between the populations of cells were a result of the inhibitors. Media with chemical inhibitors or doxycycline was replaced daily.

### CRISPR modifications of XIST constructs

Excising specific regions of the XIST construct involved transiently introducing two guide RNAs along with a Cas9 gene into HT1080 2-3-0.5a Full XIST 1c.1 cell lines. The gRNA vector pSPgRNA plasmid (Addgene #47108) gifted by Charles Gersbach contained BbsI digestible sites, that when cleaved (NEB #R0539) allowed a desired target sequence to be inserted as part of the gRNA gene. The gRNA target sequences were chosen with the E-CRISP online tool (http://www.e-crisp.org/E-CRISP/) provided by Deutsches Krebsforschungszentrum. The tools default settings were set to ‘strict’, and FASTA sequences for each region to be targeted (1kb) by a gRNA were pasted into the program. gRNAs between 22 and 19bp were included and off-target analysis was carried out using Bowtie2 against the Homo sapiens GRCh38 genome, along with an additional test for potential interference with the puromycin resistance gene.

Sense and antisense oligos of the target sequence were ordered, annealed and phosphorylated using T4 PNK (NEB #M0201). The phosphorylated double-stranded target sequences were ligated using T7 ligase (NEB #M0318) into the BbsI digested pSPgRNA plasmid and were transformed into DH5a competent cells (ThermoFisher #18265017). The pSPgRNA and the Cas9 producing plasmid pSpCas9(BB)-2A-Puro (PX459) gifted by the Zhang lab(56)(Addgene #62988) were purified using Qiaquick miniprep kits (Catalog # 27115). The concentration and purity of the purified plasmids was determined using a spectrophotometer, and in cases where the plasmids were too dilute (less than 0.5 ug/ml) they were concentrated using a speed-vac and remeasured.

The inducible XIST constructs within the HT1080 2-3-0.5a cells were modified by transiently transfecting two unique gRNA plasmids (pSPgRNA) and a Cas9 plasmid providing transient puromycin resistance (pSpCas9(BB)-2A-Puro) using Lipofectamine 3000 (ThermoFisher #L3000008). The day before the transfection the HT1080 cells were split into 24 well plates, at around ∼30-40% confluency. On the day of transfection 0.5ug of combined plasmid DNA was mixed with 1.5ul of the Lipofectamine 3000 reagent as directed by ThermoFisher’s protocol and the whole mixture was pipetted into a well of a 24 well plate. Extensive optimization determined that a molar ratio of three of each gRNA plasmid per Cas9 plasmid in the transfection mixture resulted in the greatest overall efficiency. The cells were left overnight to absorb the plasmids and 24 hours after transfection they were treated with 1ug/ml puromycin in the media.

Treatment in puromycin continued for three days at which point only cells that had undergone transfection with the pSpCas9(BB)-2A-Puro plasmid were alive. The remaining cells were then transferred to 100mm plates (roughly 20-30 per plate) and the individual cells were allowed to grow into colonies over the next two weeks. Single cell colonies were picked and transferred to 24 well plates, where they were allowed to grow until nearly confluent. When nearly confluent, the cells were split, with roughly nine tenths of the cells transferred into 1.5ml eppendorf tubes and incubated at 55°C overnight in 100ul of Mouse Homogenization Buffer (50 mM KCl, 10mM Tris-HCl, 2mM MgCl2, 0.1mg/ml gelatin, 0.45% IGEPAL CA-630, 0.45% Tween 20) with 1.2ul of 10mg/ml Proteinase K (Protocol provided by Andrea Korecki of the Simpson Lab, Centre for Molecular Medicine and Therapeutics, UBC). The Proteinase K was inactivated by incubating the samples at 95°C for ten minutes. DNA from the colonies was tested by PCR using primers spanning the regions to be deleted (Supplementary Table 2), and running the products on a gel. The colonies that produced bands of the correct size were sent to UBC’s Sequencing + Bioinformatics Consortium for Sanger sequencing. To avoid the possibility of clones arising from a single progenitor, clones that were not from different transfections needed to have at least one base pair variable at their deletion sites to be deemed independent.

### RNA isolation, reverse transcription and quantitative real-time PCR (RT-qPCR) of cDNA

RNA was obtained from cells either by directly treating cells with TRIZOL (Invitrogen) in t25 flasks or by trypsinizing and collecting adherent cells, pelleting them at 5000rcf for one minute and removing the supernatant before treating with TRIZOL. Both techniques were effective at obtaining high quality RNA and no observable difference between the results using either technique was observed. Purification of the cellular RNA was carried out according to the Invitrogen protocol in the TRIZOL user guide. The RNA obtained in this manner was measured using spectrometry to determine the concentration and purity of RNA. 5ug of RNA was transferred into 50ul DNAse1 (Roche) reactions consisting of 5ul 10x DNAse1 buffer, 1 U/ul RNAseI, 10 units of DNAse1 and the remaining volume of DEPC ddH2O. The DNAse1 reactions were incubated at 35°C for 20 minutes and then heat inactivated at 75°C for 10 minutes. The DNAse-treated RNA was then reverse transcribed using the M-MLV Reverse Transcriptase (Invitrogen #28025013) in a reaction volume consisting of 1ug of RNA, 4ul of 5x first strand buffer, 0.25mM dNTP, 0.01mM DTT, 1ul random hexamers, 1U/ul RNAse Inhibitor and 1ul of M-MLV. The final volume of 20ul was obtained with DEPC treated ddH_2_O and the reaction volume was gently mixed by inversion before letting sit at room temperature for five minutes. The reaction mixture was then incubated at 42°C for 2 hours, then inactivated at 95°C for five minutes. The newly produced cDNA was then immediately stored at −20°C until it was ready to be used in subsequent tests.

To test the relative expression of XIST across cell lines quantitative real-time PCR (qPCR) was carried out using EVAgreen (Biotium) and Maxima Hot Start Taq (Thermo Scientific) on a StepOnePlusTM Real-Time PCR System (Applied Biosystems, Darmstadt, Germany). At least three technical replicates were run for each combination of cDNA and primers tested. The cDNA itself for each sample was diluted two-fold with ddH_2_O, and the cDNA for each qPCR reaction consisted of 1.5ul of that communal dilution. Each qPCR reaction was performed in 20ul reaction volume consisting of 0.2mM dNTP, 5mM MgCl_2_, 0.25 nmoles of sense and antisense primers, 2ul of 10x Maxima Hot Start Taq Buffer, 0.16ul of Maxima Hot Start Taq and 1ul of EVA green. The remaining 18.5ul of the reaction mixture (not including the remaining 1.5ul of cDNA) consisted of ddH_2_O. For the sake of improved consistency, master mixes for each combination of primers were made and 18.5ul aliquots of that mastermix were placed in each reaction well prior to the addition of the cDNA. The PCR was performed in MicroAmp Fast Optical 96-Well Reaction Plates or Optical 8-Well Strips (Applied Biosystems) by first heating to 95°C for 5 minutes before running through 40 cycles of 95°C for 15 seconds, 60°C for 30 seconds and finally 72°C for one minute. The levels of XIST expressed in each cell line were compared to an endogenous control, PGK1, and the same primers for both genes were used across all tests and the XIST primers bound the transcript 5’ of the sites deleted. Expression of XIST in each construct was compared to a 5ddox’d Full length XIST construct whose RNA was purified and reverse transcribed simultaneously with the construct to be tested. The relative quantification of XIST in each sample was calculated using the ΔΔCT method where ΔΔCT = (CT_XIST_test_ - CT_PGK1_test_)/(CT_XIST_control_ - CT_PGK1_control_) and expressed as rq = 2^-ΔΔCt^.

### Immunofluorescence and RNA FISH

Adherent cells were transferred onto glass coverslips two days before being permeabilized and fixed, at which point growth media was removed, then the cells were washed in PBS and then in CSK buffer for 3-5 minutes. The cells were then permeabilized in chilled CSK media supplemented with Triton-X (5%v/v) for 8 minutes. The newly permeable cells were then treated in 4% paraformaldehyde in PBS for an additional 8 minutes before being stored in 70% ethanol at 4°C.

The fluorescent RNA probes for FISH were created from template DNA complementary to the XIST RNA. The probes either targeted the 5’ region of XIST including the A and F repeats, or the 3’ most 3.6kb region of the short XIST isoform’s cDNA. These probes were nick-translated (Abbott Molecular) with either Green 496 dUTP or Red 598 dUTP (Enzo) then precipitated and resuspended in 50ul of DEPC ddH_2_O and stored at −20°C in the dark.

IF and RNA FISH were performed jointly as most investigations hinged on the relative positions of various factors to the XIST RNA cloud. Permeable and fixed cells on coverslips were incubated face down on 100ul droplets of PBT (1% v/v Bovine Serum Albumin and 0.1% v/v Tween-20 in PBS) supplemented with 0.4U/ul RNAse Inhibitor (Ribolock) for twenty minutes. These cells were then incubated at room temperature for 4-6 hours in an RNAse inhibited 100ul PBT droplet, containing 1ul of the primary antibody of interest. To prevent the coverslip and antibody solution from drying they were sealed between two layers of Parafilm, creating a semi-air tight Parafilm pocket. The coverslips could also be left overnight in this Parafilm pocket without any significant repercussions, so long as they were kept at 4°C rather than room temperature.

Just prior to retrieving the coverslips from their Parafilm pockets, hybridization mixtures were created from 5ul of the nick translated fluorescent RNA probe mixed with 10ul of human Cot-1 (ThermoFisher #15279011) and 2ul of Salmon Sperm DNA (ThermoFisher #15632011), the latter two ingredients being included to prevent the probe from binding non-specifically. The hybridization mixtures were then completely dried in a speedvac while the coverslips were retrieved from their Parafilm pockets with the primary antibody solution and washed three times in PBST (PBS and 0.1% Tween-20) before being incubated for 1 hour at room temperature with the fluorescently-labelled secondary antibody (1:100 PBT supplemented with RNase Inhibitor, 1ul secondary antibody). From this point onwards the coverslips were kept in dark containers to avoid photobleaching. Next, the coverslips were washed for five minutes in PBST three times, to remove the unbound secondary antibodies. They were then fixed in 4% paraformaldehyde for 10 minutes. The coverslips were washed once in PBS to remove most of the paraformaldehyde and then were submerged for two minutes each in 70%, 80% and finally 100% ethanol to dehydrate them, before air drying the coverslips at room temperature for ∼10 minutes. The desiccated probes were resuspended in 10ul of deionized formamide (Sigma) and heated to 80°C for 10 minutes. An additional 10ul of hybridization buffer (20% BSA and 20% Dextran Sulfate in 4x SSC) was added to the hybridization mixture, which was gently mixed and pipetted as a droplet on a piece of parafilm onto which a coverslip was placed, cell side down. A second piece of parafilm was placed on top and the edges were sealed, creating a parafilm pocket where the probes could hybridize overnight at 37°C.

The next day the coverslips were retrieved and incubated in an equal measure of deionized formamide (Sigma) and 4x SSC (Invitrogen) at 37°C for 20 minutes. The coverslips were then incubated in 2x SSC at 37°C and 1x SSC at room temperature, for 15 minutes each. DAPI staining was performed by placing the coverslips in a 0.1ug/ml solution of DAPI in pure methanol at 37°C for 15 minutes. The excess DAPI was rinsed off in methanol and the coverslips were mounted on glass slides using a hardset antifade mounting media (Vectashield). After letting the media harden, cells were photographed using a confocal fluorescence microscope (DMI 6000B) and camera (MicroPublisher 5.0 RTV, Qimaging) to capture and compile the red (secondary antibody), green (XIST RNA probe) and blue (DAPI) fluorescent channels using the Openlab program (Perkin Elmer).

The colour channel images obtained for a given cell were converted into composite RGB images using ImageJ (Fiji). The fluorescent intensity of the DAPI (blue), immunofluorescence channel (red), and XIST fluorescence (green), were measured using the BAR plugin and drawing a straight line bisecting the point of greatest XIST fluorescence and the maximal width of the nucleus possible without intersecting a nucleolar territory. The fluorescent intensity at each position across the line was recorded and the positions were bifurcated into either the XIST +ve category or XIST -ve category. The pixels that had a level of green intensity greater than 50% of the maximum along the range of XIST fluorescence were defined as XIST +ve while those less than 25% were defined as XIST negative. Those very few positions with XIST signal intensity between 25%-50% were not included in the analysis to better delineate the two groups. The mean and standard deviation of Immunofluorescence intensity in the XIST +ve and XIST -ve groups were determined and used to calculate the z-score for each cell. Either 59-61 cells were analyzed in every condition described throughout this work and the population distribution of z-scores were compared using a Mann-Whitney U test. The threshold for statistical significance was adjusted based on the number of tests being performed (e.g. p-value = 0.05/# of tests). The identities of all the cells tested and the heterochromatin marks being analyzed in each case were blinded throughout this process and the identities only revealed after all testing and analysis had been completed.

### Western Blotting

Protein was extracted from the HT1080 cell lines using RIPA buffer, prepared according to the Cold Spring Harbor protocol, supplemented with 1µl of Protease Inhibitor Cocktail from Roche per 100µl of RIPA buffer. A volume of 500µl of RIPA buffer was used per 2 million cells used. Samples were gently agitated by shaking for 30 minutes at 4°C to allow for the breakdown of cells, then were centrifuged at max speed (>14,000 rcf) at 4°C for 20 minutes.

The acrylamide gel was prepared in Bio-Rad 1.5mm casting plates and gel casting stand. The lower gel consisted of 10ml of 12% acrylamide lower running gel [4.2ml of 29:1 acrylamide/bis-acylamide (Bio Rad), 2.5ml of 4x lower gel buffer (1.5M Tris and 0.4%SDS in distilled water, pH of 8), 3.3ml of distilled water, 10µl of TEMED from Fisher Scientific and 40µl of 10% ammonium persulfate (which was prepared fresh each time) and topped with 100% isopropyl alcohol ∼2cm from the top of the plates. Once the lower gel solidified in the plates, the upper gel mixture replaced the isopropyl alcohol [0.75ml 29:1 acrylamide/bis-acylamide, 0.45ml of 4x upper gel buffer (0.5M Tris and 0.4%SDS in distilled water, pH of 6.8), 1.8ml of distilled water, 10µl of of 10% ammonium persulfate and 5µl of TEMED (Fisher Scientific)] and a gel comb was inserted. The solidified gel was loaded into a Bio-Rad electrode assembly and buffer tank. The 1x running buffer was diluted from a 10x stock (0.25M Tris, 1.92M Glycine and 1% w/v SDS in distilled water) using distilled water. 15µl of each protein extract was mixed with 2X SDS gel-loading buffer (4% SDS, 0.2% bromophenol blue, 20% glycerol and 200mM dithiothreitol) according to the Cold Spring Harbor protocol and the resulting mixture was heated to 95°C for 2-3 minutes. The samples were run with 180 volts until the leading band reached the bottom of the gel. BenchMark Pre-stained Protein Ladder from Thermo Fisher was loaded into one of the wells abutting the samples to provide a reference for the size of the bands.

To transfer the protein samples to a piece of 0.2µm nitrocellulose paper (ThermoFisher) the gel and nitrocellulose paper were placed together inside a Bio Rad Core Assembly Module according to the manufacturer’s instructions. The Core Assembly Module was then inserted as directed into the Bio Rad protein transfer tank and a cold ice pack was placed in the Transfer Tank as well. The tank was then filled with Transfer Buffer diluted from a cold 10x stock [0.25M Tris and 1.9M glycine] with 20% methanol and 70% distilled water per final volume. and was hooked up to a Bio Rad PowerPac HC power supply. Protein transfer was carried out at 90 volts for one hour at 4°C with constant gentle agitation of the Transfer buffer using a magnetic stir bar to ensure dispersion of heat.

After the proteins were transferred to the nitrocellulose membrane the membrane was incubated with a 40ml blocking buffer (0.1% v/v Tween-20 and 3% Bovine Serum Albumin) for 1 hour while gently being agitated to ensure complete coverage of the buffer. After blocking, the nitrocellulose was placed inside a 50ml falcon tube containing 20ml of primary antibody solution (3% w/v BSA, 0.1% v/v Tween-20 in TBS plus the relevant antibodies). The membrane was incubated in this solution overnight at 4°C on a tube rotator to allow even coating of the entire surface of the membrane with antibodies. The next day the nitrocellulose membrane was washed four times for 5 minutes each with TBST (0.1% v/v Tween-20 in TBS). The nitrocellulose membrane was then in 20ml of secondary antibody solution (3% w/v BSA, 0.1% v/v Tween-20 in TBS plus either goat anti-mouse or goat anti-rabbit fluorescently labelled secondary antibodies) for 1 hour while being gently agitated. The nitrocellulose was washed twice with TBST then once with TBS for five minutes each. The protein and antibody bound membrane was then imaged at the relevant wavelengths using an LI-COR Odyssey machine from BioAgilytix and the software package, Image Studio.

## Acknowledgements

We would like to thank Bradley Balaton for his help with the computational analysis of the enrichment z-score calculations. The Brown lab research group of Sarah Baldry, Christine Yang, Sam Peeters and Kira Tosefsky are also thanked for their advice throughout this project.

## Author Contributions

TDM^c^D performed all of the experiments, data analysis, designed the figures and tables and contributed to planning the research and writing the manuscript. CJB supervised the project and contributed to planning the project and writing the manuscript.

## Supplementary Material

**Supplementary Figure 1:**
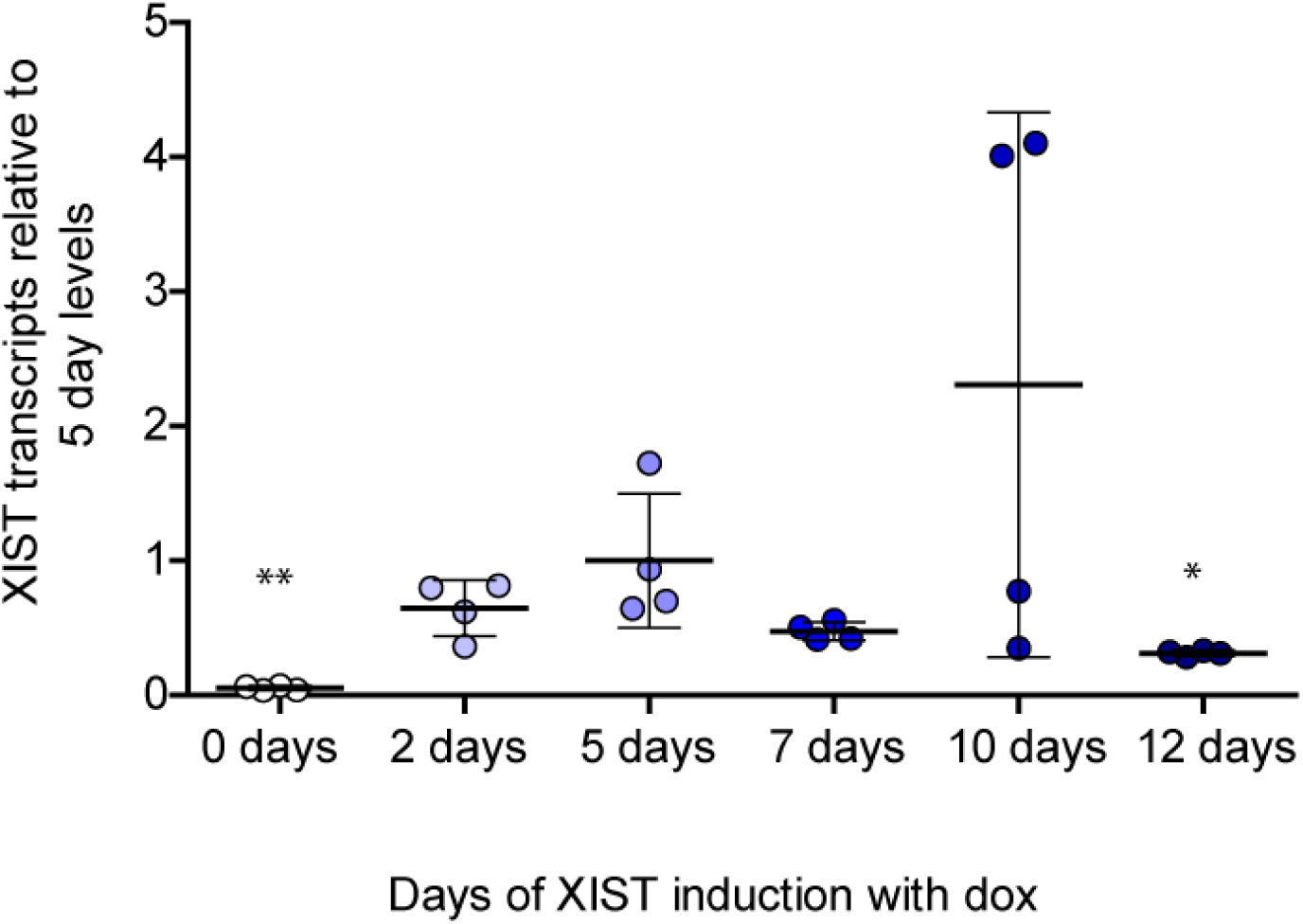
Relative XIST transcript levels expressed from chromosome 8p in HT1080 cell line. *XIST* transcript levels were measured using RT-qPCR and determining the relative transcript levels compared to the endogenous control gene *PGK1*. Four biological replicates for each time point of *XIST* induction in the 8p HT1080 cell line were tested and the relative levels of *XIST* were normalized to the average level of the 5 day time point, as it was the time point which had become standard in previous examinations of the model system. The mean and standard deviation for each condition are indicated by lines, and individual dots representing the relative expression of each replicate. Increasing darkness of shading was used to indicate increasing length of *XIST* induction. The statistical significance of a difference between the 5 day treatment and other time points was calculated by two-tailed unpaired t-test (* p < 0.05, ** p< 0.01).

**Supplementary Table 1:**
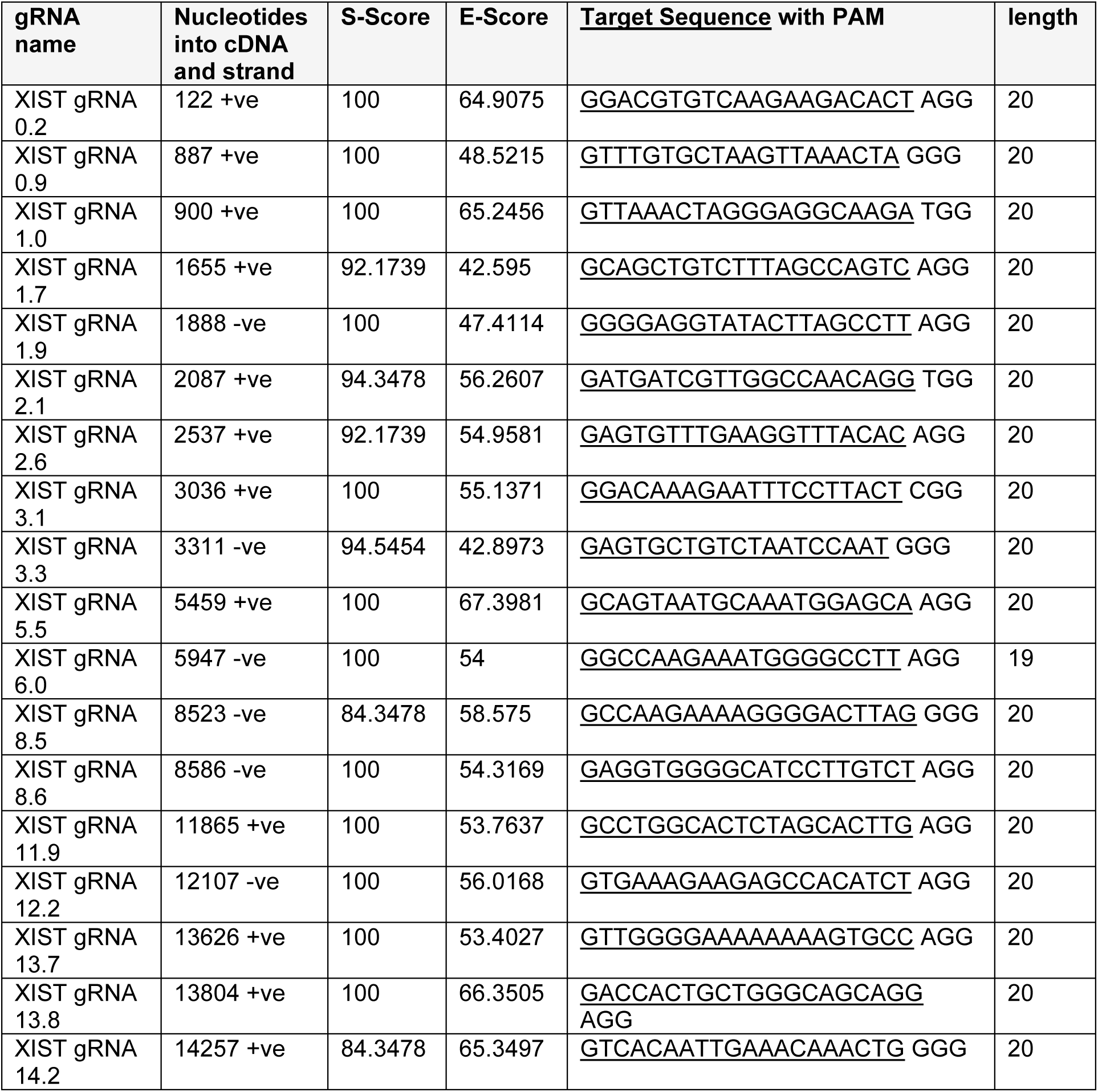
gRNAs used to generate XIST deletions

**Supplementary Table 2:** XIST deletion sizes confirmed by sequencing across deletion. Each of the cell lines successfully generated for each type of deletion construct are listed along with the gRNAs used to create the deletion and the total number of nucleotides lost from the XIST cDNA sequence.

| Deletion cell line | 5' gRNA | 3' gRNA | Nucleotides deleted |
| --- | --- | --- | --- |
| Δ A #12 | *XIST* gRNA 0.2 | *XIST* gRNA 0.9 | 777 |
| Δ FBh #21 | *XIST* gRNA 0.9 | *XIST* gRNA 1.9 | 1127 |
| Δ FBh #22 | *XIST* gRNA 1.0 | *XIST* gRNA 1.9 | 811 |
| Δ Bh #5 | *XIST* gRNA 1.7 | *XIST* gRNA 2.6 | 833 |
| Δ Bh #7 | *XIST* gRNA 1.7 | *XIST* gRNA 2.6 | 833 |
| Δ Bh #11 | *XIST* gRNA 1.7 | *XIST* gRNA 2.6 | 857 |
| Δ BC #2 | *XIST* gRNA 2.1 | *XIST* gRNA 3.3 | 1195 |
| Δ BC #8 | *XIST* gRNA 2.1 | *XIST* gRNA 3.3 | 1189 |
| Δ BC #17 | *XIST* gRNA 2.1 | *XIST* gRNA 3.3 | 1195 |
| Δ 3'PfIMI #3 | *XIST* gRNA 3.1 | *XIST* gRNA 6.0 | 2859 |
| Δ 3'PfIMI #6 | *XIST* gRNA 3.1 | *XIST* gRNA 6.0 | 2859 |
| Δ D #3 | *XIST* gRNA 5.5 | *XIST* gRNA 8.5 | 3084 |
| Δ D #10 | *XIST* gRNA 5.5 | *XIST* gRNA 8.5 | 3092 |
| Δ 3D5E #13 | *XIST* gRNA 8.5 | *XIST* gRNA 12.2 | 3584 |
| Δ 3D5E #14 | *XIST* gRNA 8.5 | *XIST* gRNA 12.2 | 3583 |
| Δ 3D5E #15 | *XIST* gRNA 8.5 | *XIST* gRNA 12.2 | 3588 |
| Δ E #6 | *XIST* gRNA 11.9 | *XIST* gRNA 13.7 | 1844 |
| Δ E #10 | *XIST* gRNA 11.9 | *XIST* gRNA 13.7 | 1778 |
| Δ 3' #1 | *XIST* gRNA 13.7 | *XIST* gRNA 14.2 | 630 |
| Δ 3' #7 | *XIST* gRNA 13.7 | *XIST* gRNA 14.2 | 630 |

**Supplementary Table 3:** XIST expression levels in deletion clones relative to Full XIST. The relative expression (RQ) of each deletion cell line as well as construct is listed along with the standard deviation (SD) across biological replicates (≥3 per cell line) as well as the resulting p value of any construct or cell lines difference from Full XIST.

| Construct | Clone | Mean RQ | SD | p-value | Mean | SD | p-value |
| --- | --- | --- | --- | --- | --- | --- | --- |
| $\Delta A$ | 12 | 0.508 | 0.300 | 1.9E-02 | 0.508 | 0.300 | 1.90E-02 |
| $\Delta FBh$ | 21 | 0.678 | 0.425 | 9.6E-02 | 1.044 | 0.686 | 5.71E-01 |
|  | 22 | 1.410 | 0.764 | 4.2E-01 |  |  |  |
| $\Delta Bh$ | 5 | 0.971 | 0.319 | 5.3E-01 | 0.813 | 0.315 | 1.01E-01 |
|  | 7 | 0.527 | 0.232 | 4.8E-02 |  |  |  |
|  | 11 | 0.942 | 0.184 | 4.3E-01 |  |  |  |
| $\Delta PfIMI$ | 3 | 0.630 | 0.171 | 9.0E-02 | 0.630 | 0.171 | 8.96E-02 |
| $\Delta BC$ | 2 | 0.653 | 0.235 | 3.1E-02 | 0.773 | 0.272 | 1.71E-02 |
|  | 8 | 0.905 | 0.328 | 3.7E-01 |  |  |  |
|  | 17 | 0.762 | 0.166 | 1.2E-01 |  |  |  |
| $\Delta 3'PfIMI$ | 3 | 1.147 | 0.460 | 9.1E-01 | 1.150 | 0.489 | 8.88E-01 |
|  | 6 | 1.154 | 0.526 | 9.1E-01 |  |  |  |
| $\Delta D$ | 2 | 1.258 | 0.919 | 6.8E-01 | 1.061 | 0.759 | 9.02E-01 |
|  | 10 | 0.864 | 0.360 | 3.0E-01 |  |  |  |
| $\Delta 3D5E$ | 13 | 1.249 | 0.919 | 7.0E-01 | 1.075 | 0.857 | 8.31E-01 |
|  | 14 | 1.367 | 1.059 | 5.6E-01 |  |  |  |
|  | 15 | 0.610 | 0.128 | 3.3E-02 |  |  |  |
| Exon 1 | 3 | 1.072 | 0.043 | 8.5E-01 | 1.321 | 0.655 | 4.66E-01 |
|  | 7 | 1.571 | 0.855 | 2.4E-01 |  |  |  |
| $\Delta E$ | 6 | 1.762 | 1.085 | 8.1E-02 | 1.797 | 1.017 | 4.02E-02 |
|  | 10 | 1.833 | 0.930 | 3.5E-02 |  |  |  |
| $\Delta 3'$ | 1 | 0.365 | 0.115 | <u>1.1E-03</u> | 0.395 | 0.169 | <u>2.04E-05</u> |
|  | 7 | 0.425 | 0.199 | <u>1.0E-03</u> |  |  |  |
| $\Delta\Delta$ | 12 | 0.986 | 0.739 | 6.4E-01 | 0.986 | 0.739 | 6.40E-01 |

**Supplementary Table 4:** Summary of H3K27me3 enrichment in deletion constructs. List of the number of cells analyzed, the median z-score calculated as well as the standard deviation (SD) for each construct. The statistical significance of each population of deletion constructs’ difference from Full XIST in their enrichment was calculated using the Mann-Whitney U test and the p values are listed.

| H3K27me3 | number of cells | median z-score | SD | MW p-value |
| --- | --- | --- | --- | --- |
| Full *XIST* | 60 | 2.590 | 1.704 | -- |
| Δ A | 60 | 1.590 | 1.196 | <u>1.16E-04</u> |
| Δ FBh | 59 | -0.147 | 1.568 | <u>1.11E-13</u> |
| Δ Bh | 59 | 0.787 | 4.869 | <u>8.47E-07</u> |
| Δ PflMI | 60 | 1.990 | 1.181 | 1.11E-02 |
| Δ BC | 60 | 1.910 | 1.127 | 1.17E-03 |
| Δ 3'PflMI | 61 | 2.040 | 1.248 | 1.54E-02 |
| Δ D | 60 | 1.900 | 1.519 | 1.55E-02 |
| Δ 3D5E | 61 | 2.110 | 1.912 | 1.01E-01 |
| Exon 1 | 60 | 0.110 | 1.407 | <u>1.23E-14</u> |
| Δ E | 60 | -0.010 | 0.860 | <u>2.97E-17</u> |
| Δ 3' | 60 | 2.140 | 1.266 | 1.99E-02 |
| ΔΔ | 60 | 0.338 | 1.252 | <u>2.42E-15</u> |

**Supplementary Table 5:** Summary of ubH2A enrichment in deletion constructs

| UbH2A | number of cells | median z-score | sd | M.W. p-value |
| --- | --- | --- | --- | --- |
| Full *X/ST* | 60 | 4.178 | 3.179 | -- |
| Δ A | 60 | 0.223 | 1.628 | <u>2.84E-17</u> |
| Δ FBh | 60 | 3.328 | 2.937 | 1.48E-01 |
| Δ Bh | 60 | 5.972 | 4.801 | 6.30E-03 |
| Δ PflMI | 60 | -0.180 | 1.776 | <u>1.83E-16</u> |
| Δ BC | 60 | -0.653 | 0.712 | <u>4.35E-21</u> |
| Δ 3'PflMI | 61 | 2.848 | 3.237 | 3.65E-02 |
| Δ D | 60 | -0.425 | 0.694 | <u>8.30E-21</u> |
| Δ 3D5E | 60 | 3.052 | 2.987 | 8.95E-02 |
| Exon 1 | 59 | -0.352 | 2.896 | <u>1.20E-16</u> |
| Δ E | 60 | 2.523 | 3.184 | 5.91E-03 |
| Δ 3' | 60 | -0.635 | 0.714 | <u>4.80E-21</u> |
| ΔΔ | 58 | 0.039 | 1.696 | <u>7.71E-17</u> |

**Supplementary Table 6:** Summary of MacroH2A enrichment in deletion constructs

| MacroH2A | number of cells | median z-score | sd | MW p-value |
| --- | --- | --- | --- | --- |
| Full *X/ST* | 60 | 1.958 | 1.469 | -- |
| Δ A | 58 | 2.213 | 2.227 | 6.26E-01 |
| Δ FBh | 59 | -0.434 | 1.189 | <u>5.34E-17</u> |
| Δ Bh | 60 | 1.709 | 2.639 | 2.90E-01 |
| Δ PflMI | 60 | -0.348 | 1.124 | <u>1.89E-17</u> |
| Δ BC | 60 | 1.562 | 2.247 | 8.62E-03 |
| Δ 3'PflMI | 60 | 0.313 | 2.197 | <u>3.06E-09</u> |
| Δ D | 60 | -1.042 | 2.338 | <u>4.25E-17</u> |
| Δ 3D5E | 60 | 1.541 | 1.444 | 1.09E-01 |
| Exon 1 | 59 | -0.096 | 1.380 | <u>9.44E-15</u> |
| Δ E | 60 | -1.173 | 1.784 | <u>6.08E-17</u> |
| Δ 3' | 60 | 2.609 | 2.288 | 7.58E-02 |
| ΔΔ | 61 | -0.737 | 1.731 | <u>3.76E-18</u> |

**Supplementary Table 7:** Summary of SMCHD1 enrichment in deletion constructs

| SMCHD1 | number of cells | median z-score | sd | MW p-value |
| --- | --- | --- | --- | --- |
| Full *X/ST* | 60 | 2.704 | 2.167 | -- |
| Δ A | 60 | 2.601 | 2.307 | 4.36E-01 |
| Δ FBh | 60 | 2.161 | 1.521 | 5.00E-02 |
| Δ Bh | 60 | 2.249 | 1.576 | 6.13E-02 |
| Δ PflMI | 60 | -0.109 | 1.179 | <u>1.04E-14</u> |
| Δ BC | 60 | -0.516 | 1.073 | <u>1.54E-16</u> |
| Δ 3'PflMI | 61 | 2.902 | 1.951 | 9.98E-01 |
| Δ D | 60 | -0.350 | 1.629 | <u>2.47E-14</u> |
| Δ 3D5E | 60 | -0.386 | 1.011 | <u>6.35E-17</u> |
| Exon 1 | 60 | -0.407 | 1.003 | <u>1.05E-17</u> |
| Δ E | 61 | 3.301 | 2.814 | 1.89E-01 |
| Δ 3' | 60 | 1.291 | 1.582 | <u>2.01E-07</u> |
| ΔΔ | 60 | -0.283 | 1.989 | <u>2.64E-15</u> |

**Supplementary Table 8:** Summary of effect of chemical inhibitors on XIST mediated chromatin remodelling. List of the number of cells analyzed, the median z-score calculated as well as the standard deviation (SD) for each treatment condition and heterochromatin feature. The statistical significance of each population of inhibitor treated cells’ difference from the uninhibited control population was calculated using the Mann-Whitney U test and the p values are listed.

| Treatment | Mark | number of cells | median z-score | sd | MW p-value |
| --- | --- | --- | --- | --- | --- |
| Control | H3K27me3 | 60 | 2.593 | 1.704 | -- |
| GSK343 5uM | H3K27me3 | 58 | -0.090 | 1.479 | 7.24E-16 |
| PRT4165 50uM | H3K27me3 | 60 | 2.483 | 1.648 | 7.59E-01 |
| Control | UbH2A | 60 | 4.178 | 3.179 | -- |
| GSK343 5uM | UbH2A | 60 | 3.053 | 5.754 | 2.18E-01 |
| PRT4165 50uM | UbH2A | 60 | 0.469 | 0.846 | 5.28E-18 |
| Control | MacroH2A | 60 | 1.958 | 1.469 | -- |
| GSK343 5uM | MacroH2A | 60 | 0.224 | 1.231 | 1.30E-10 |
| Control | SMCHD1 | 60 | 2.704 | 2.167 | -- |
| PRT4165 50uM | SMCHD1 | 60 | -0.196 | 1.215 | 5.28E-18 |

